# Sequential coupling of sleep oscillations enables concept-neuron reactivation and supports information flow across the human hippocampal-cortical circuit

**DOI:** 10.64898/2026.01.15.699122

**Authors:** Fabian Schwimmbeck, Johannes Niediek, Valeri Borger, Rainer Surges, Eugen Trinka, Thomas Schreiner, Florian Mormann

## Abstract

Effective memory consolidation during sleep is thought to rely on the transfer of reactivated memory traces from the hippocampus to the cortex. However, the mechanisms supporting this essential dialogue across brain areas remain poorly understood. Here, we recorded single-neuron activity (n = 1097) in the medial temporal lobe (MTL) of 10 epilepsy patients across 21 nights of sleep. Before sleep, patients engaged in a memory task with stimuli eliciting concept-neuron responses. During non-rapid eye movement (NREM) sleep, neuronal firing locked to sharp wave–ripples (SWR) revealed a directed flow of information from the hippocampus to cortical targets. Notably, within SWR-driven activity, experience-dependent concept neurons exhibited elevated co-activation both within and across MTL regions, indicating selective reactivation of behaviorally relevant information. Cross-regional co-activation of concept cells was markedly enhanced when hippocampal SWRs coincided with cortical slow oscillation–spindle complexes, suggesting an active role of the cortex in shaping interregional communication. These findings provide evidence that SWRs in humans selectively reactivate experience-related neural ensembles across the hippocampal–cortical network, while synergistic interactions with cortical slow oscillation–spindle events might facilitate effective memory consolidation.

## Introduction

The consolidation of episodic memories is thought to rely on sleep-related neural processes that stabilize and integrate newly encoded experiences (Buzsáki, 1989; Diekelmann & Born, 2010a). However, the underlying mechanisms remain poorly understood in humans, largely due to the involvement of deep MTL structures that lie beyond the reach of non-invasive human imaging techniques (Parvizi & Kastner, 2018).

Invasive studies in rodents have shown that neurons in the MTL encode the spatiotemporal structure of experienced environments, forming behaviorally relevant activity patterns (Moser et al., 2008; O’Keefe, 1976). During subsequent NREM sleep, these patterns are reactivated in the hippocampus, typically in conjunction with SWRs (Nádasdy et al., 1999; Ólafsdóttir et al., 2016; Skaggs & McNaughton, 1996; Wilson & McNaughton, 1994) Such reactivation is thought to enable the transfer of memory representations from the hippocampus to long-term neocortical stores, stabilizing initially labile memories (Buzsáki, 1989). This essential hippocampal–cortical dialogue is assumed to be orchestrated by the coordinated interplay of SWRs with the cardinal NREM sleep-specific oscillations, namely cortical slow oscillations (SOs) and thalamo-cortical sleep spindles (Cairney et al., 2018; Schreiner et al., 2021, 2022; Schwimmbeck et al., 2025; Sirota et al., 2003; Staresina et al., 2015). While some of the key mechanisms proposed above have been characterized in animal models, studies in humans have largely relied on indirect evidence (e.g., EEG and invasive EEG, (Schönauer et al., 2017; Schreiner et al., 2021, 2024), fMRI ((Bergmann et al., 2012; Peigneux et al., 2004; Sterpenich et al., 2021)), as direct access to single-neuron activity, which is critical for evaluating these fundamental questions, remains limited. In particular, it remains unknown whether human hippocampal SWRs drive the endogenous reactivation of experience-linked neurons, and whether coordinated sleep oscillations support the exchange of memory-related information between brain regions.

Reminiscent of rodent place cells tuned to spatial locations, concept neurons in the human MTL encode abstract semantic representations of the external world (Kehl et al., 2024; Quian Quiroga et al., 2009; Quiroga et al., 2005). Because their activity can be directly linked to experience and later recall (Staresina et al., 2019), concept neurons have been hypothesized to provide the semantic building blocks of episodic memory (Quiroga, 2012), a proposal that has recently received experimental support (Mackay et al., 2024). This makes them uniquely suited for investigating the fundamental mechanisms of memory reactivation and the hippocampal–cortical dialogue underlying sleep-dependent memory consolidation. Here, we hypothesize that SWRs orchestrate a directed inter-regional flow of information, through which ensembles of experience-related concept neurons in the MTL are selectively reactivated. We further posit that their temporal coupling to cortical SO–spindle complexes strengthens interregional coactivation of single neurons, promoting hippocampal–cortical communication (Fig. 1A).

**Figure 1.**
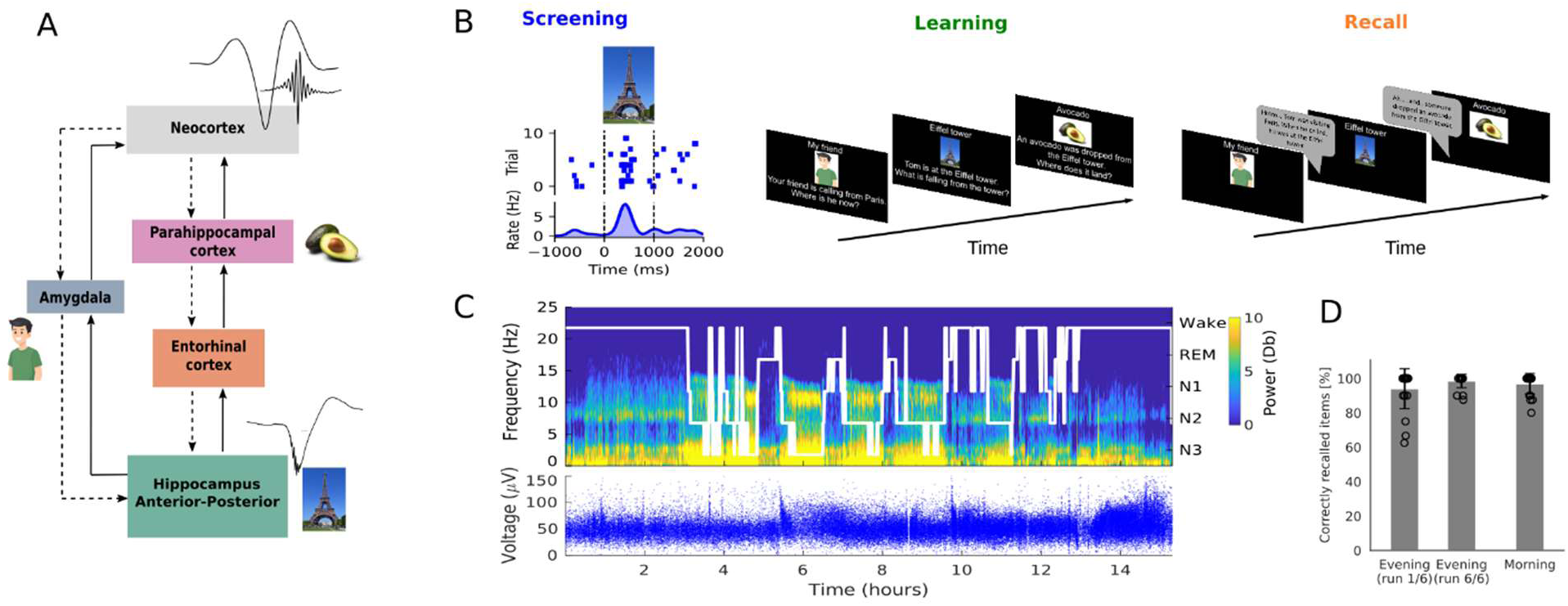
Working model, experimental procedure, and behavioral outcome. (**A**) Simplified scheme of hippocampal-cortical information flow, interconnected MTL regions, including semantically tuned neurons (e.g., Eiffel tower-neuron). Hippocampal SWRs selectively coactivate semantically tuned neurons in the hippocampus and downstream regions (SWR information flow indicated by solid arrows) during NREM sleep. Reactivation and cross-regional relaying of mnemonic information is timed and facilitated by neocortical feed-forward input during cortical SO-spindle complexes (top-down information flow, dashed lines). To examine this model, bi-temporal multisite recordings were conducted simultaneously in all depicted MTL regions alongside scalp EEG. (**B**) As detailed in Niediek et al., 2026, experimental sessions started with an initial stimulus screening to identify selectively tuned neurons. Response-eliciting stimuli were then incorporated into an individually tailored image- and text-based narrative (”Fotonovela”). Patients learned the narrative at their own pace until they felt confident to recite it correctly. Recall performance was tested six times in the evening before sleep and once in the next morning. Stability of stimulus selectivity of responsive neurons was tested with mini-screenings conducted before and after the learning session, as well as in the next morning (see also Niediek et al., 2026). (**C**) Top panel: Exemplary hypnogram and corresponding time-frequency spectrogram (0-25 Hz) across a full experimental session. Bottom panel shows recording stability of the previously identified Simpsons-neuron across time. (**D**)Recall performance approached ceiling level in both the 1st (94.0 ± 5.2%) and 6th (98.5 ± 2.7%) pre-sleep runs, and remained high in the following morning (96.9 ± 3.8%).

To address these hypotheses, we leveraged the rare opportunity to perform simultaneous single-neuron recordings from multiple sites within the human MTL, alongside intracranial EEG (macro-electrode recordings, iEEG) and scalp EEG recordings during both wakefulness and sleep. Data were collected from 10 neurosurgical patients, who performed an episodic memory task prior to sleep (Niediek et al., 2026), 21 sessions in total). We show that SWRs in the human hippocampus drive a directed flow of information toward cortical targets during NREM sleep, as indicated by progressively increasing time lags in neural firing. Specifically, within SWR-locked neural activity, experience-dependent concept neurons were selectively co-activated across the hippocampal–cortical network, indicating memory reactivation The strongest coactivation occurred when SWRs coincided with SO–spindle complexes, suggesting a synergistic role of these cortical rhythms in facilitating interregional communication for memory consolidation. By revealing how hippocampal SWRs and cortical rhythms jointly govern the reactivation of experience-related single-neuron activity, our findings critically advance the mechanistic understanding of sleep-dependent memory consolidation in the human brain.

## RESULTS

### Behavior

The behavioral task has been described in full detail elsewhere (Niediek et al., 2026). Briefly, each patient participated in 1 to 4 experimental night sessions (mean ± SEM: 2.1 ± 0.3). In each session, we first conducted a screening procedure to identify 8 to 10 stimuli (mean ± SEM: 9.2 ± 0.17) to which neurons responded with selective increases in firing. The corresponding concepts were then embedded into a personalized narrative (“Fotonovela”, Fig. 1 B), which participants were instructed to learn prior to sleep. The narrative was presented as a slideshow on a laptop, with each slide featuring a single concept shown to elicit a neural response during the screening session conducted earlier on the same day. Participants learned the narrative at their own pace, followed by six consecutive recall trials before sleep. A final recall session took place in the following morning after a full night of sleep (mean sleep duration ± SEM: 7.06 ± 0.34 h; for sleep characteristics see Table S1). The purpose of this task was to imprint robust neural representations of the learning material that could be tracked during sleep. Recall performance remained near ceiling across all sessions (final recall pre-sleep (mean ± SEM): 98.45 ± 2.96%; post-sleep: 96.9 ± 3.78%; Fig. 1D). In total, we recorded from 1097 neurons (849 non-responsive neurons, mean ± SEM: 40.33 ± 5.23; 248 stimulus selective neurons, mean ± SEM: 11.81 ± 1.32 per session) across 21 full nights of sleep in 10 neurosurgical patients. To ensure stability of stimulus selectivity throughout the experiment, brief screening sessions were conducted before learning, after the final evening recall, and again prior to the morning recall. Recordings took place at the Department of Epileptology at the University Hospital Bonn, Germany, during routine presurgical evaluation with depth electrodes implanted for invasive seizure monitoring.

### Sharp wave-ripples direct hippocampal-to-cortical information transfer

The transfer of reactivated memory content from the hippocampus to the cortex, central to systems consolidation, is thought to be initiated by SWRs (Buzsáki, 2015; Todorova & Zugaro, 2020). If SWRs drive bottom-up information flow, this should be reflected in systematic time lags of SWR-locked firing along hippocampal-cortical pathways. However, direct evidence for this directional flow of information remains elusive, largely due to the rarity of simultaneous recordings across multiple brain regions (Nitzan et al., 2022). To directly test whether hippocampal SWRs orchestrate such guided communication, we performed multisite single-neuron recordings in the MTL during human sleep. Neural activity was sampled along the hippocampal longitudinal axis (anterior [AH] and posterior [PH] segments) as well as across major hippocampal–cortical output pathways, including the entorhinal cortex (EC), parahippocampal cortex (PHC), and amygdala (A) (see Fig. 1A for illustration).

Hippocampal SWRs (80 -120 Hz in humans) were detected in the anterior hippocampus (AH) from bipolar-referenced macroelectrode contacts during NREM sleep stage 2 and slow-wave sleep (SWS), following established procedures (Helfrich et al., 2019; Staresina et al., 2015, 2023). To ensure physiological validity, segments containing interictal epileptiform discharges (IEDs) were excluded using automated detection algorithms ((Staresina et al., 2015), see Fig. S2). Across 21 sessions, we identified 50,118 AH SWR events (mean ± SEM: 1261.19 ± 136.08 per session). Detected events exhibited a mean frequency of 92.27 ± 0.04 Hz (SEM), a mean duration of 56 ± 0.08 ms (SEM), and a mean amplitude of 10.09 ± 0.02 µV (SEM) (see Fig. 2A and B), consistent with previously reported characteristics of hippocampal SWRs in humans (Helfrich et al., 2019; Kunz et al., 2024; Liu et al., 2022; Staresina et al., 2015, 2023). To assess whether AH detected SWRs would drive neural activation from the hippocampus to cortical regions in a stepwise manner, we examined SWR-locked firing patterns across hippocampal subregions (i.e., AH, PH) and distributed MTL areas (i.e., EC, PHC, and A). For each session, hemisphere, region, and neuron, we created peri-event time histograms (PETHs) locked to SWR peaks (±500 ms). PETHs were normalized using a 1,000 ms baseline period preceding the event (-1500 to -500 ms relative to SWR peaks). Peak firing times were identified within ±75 ms around SWR peaks (Fig. 2C). Finally, we compared the distributions of these peak firing time lags along the hippocampal longitudinal axis (anterior to posterior) and across the hippocampal–cortical output pathway (Fig. 2D).

**Figure 2.**
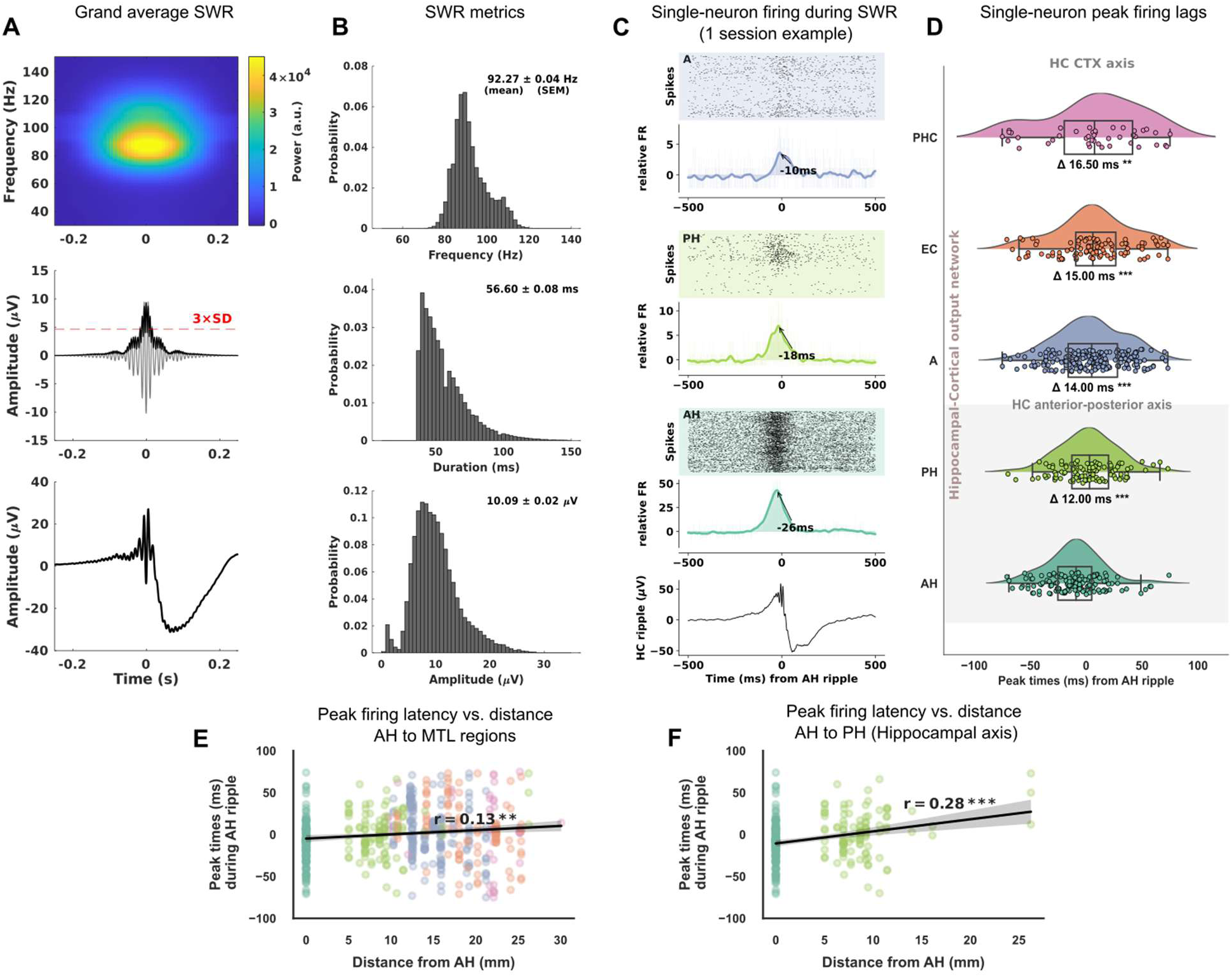
SWRs direct hippocampal-to-cortical information flow. (**A**) SWRs detected in the anterior hippocampus (n = 50,118; mean ± SEM: 1261.19 ± 136.08 per session). *Top*: Averaged time–frequency representation; *middle*: SWR-aligned event-related potential (ERP) filtered between 80–120 Hz, including RMS signal and detection threshold (3 SDs above mean of RMS signal) for candidate events, with SWR duration defined by the time points at which the signal crossed this threshold; *bottom*: corresponding unfiltered grand average SWR trace of hippocampal iEEG, centered on individual SWR peaks in the RMS signal. (**B**) Summary of SWR characteristics across sessions, including frequency distribution (mean ± SEM: 92.27 ± 0.04 Hz), duration (mean ± SEM: 56.6 ± 0.08 ms), and amplitude (mean± SEM: 10.09 ± 0.02 µV) (**C**) Example from a single session showing single-neuron activity time-locked to anterior hippocampal (AH) SWRs. Colored raster plots and smoothed spike-count histograms depict SWR-associated firing across regions, with time lags defined by the peak firing latency within ±75 ms of the SWR peak. (**D**) Group-level distribution of regional peak firing latencies across 21 sessions. Δ time lags represent the median delay of SWR-locked firing bursts in each region relative to SWR-locked anterior hippocampal neurons. Positive values indicate delayed responses downstream of the AH seed region. Box plots depict the median and interquartile range. *P*-values are derived from two-sided Wilcoxon rank-sum test against AH time lags, with Holm-correction for multiple comparisons. **(E)** Neural peak firing latency during AH SWRs is significantly correlated with distance from the anterior hippocampus across MTL regions; **(F)** along the hippocampal anterior–posterior axis. \*\*\**p* < 0.001, \*\**p* < 0.01. Anterior hippocampus (AH), posterior hippocampus (PH), entorhinal cortex (EC), parahippocampus (PHC), amygdala (A), cortex (CTX).

Supporting the notion of SWR-initiated information flow, we observed systematic delays in peak firing times along both the hippocampal longitudinal axis and the hippocampal-cortical output pathway. Compared to SWR-locked bursts in the AH, median peak firing in the PH was significantly delayed by 12 ms (*p* = 2.57 × 10^-4^) Extending to downstream cortical targets (i.e., EC, PHC, and A), we found a graded increase in peak firing lags relative to AH detected SWRs (A: 14 ms, *p* = 1.3 × 10^−5^, EC: 15 ms, *p* = 1.3 × 10^−5^; PHC: 16.5 ms, *p* = 1.12 × 10^-3^; all Wilcoxon rank-sum tests, Holm-corrected). Notably, a similar SWR-triggered propagation pattern, consistent with AH serving as a functional origin, was evident for PH detected SWRs (Fig. S3).

To further characterize the spatial organization of SWR-locked firing delays, we examined the relationship between peak firing latency and anatomical distance from the anterior hippocampus. Across MTL regions, peak firing latencies during AH SWRs exhibited a significant positive correlation with distance from AH, such that neurons located at greater distances fired progressively later relative to AH SWR peaks (Fig. 2E; Pearson *r* = 0.13, *p* = 1.61 × 10^−3^). An analogous distance–latency relationship was observed within the hippocampus itself, with firing latencies increasing systematically along the anterior–posterior axis away from AH (Fig. 2F; Pearson *r* = 0.28, *p* = 7.6 × 10^−6^). Together, these results provide direct evidence that SWRs orchestrate directed, cross-structural neuronal communication at single-neuron resolution during NREM sleep, with the anterior hippocampus acting as a principal source of coordinated information flow across hippocampal and hippocampal–cortical pathways.

### Coordinated sleep oscillations enable bidirectional hippocampal–cortical communication

Endogenous communication from the hippocampus to the cortex is thought to be coordinated by sequential slow oscillation–spindle complexes, organizing large-scale network dynamics to align hippocampal signaling with periods of heightened cortical receptivity (Isomura et al., 2006; Niethard et al., 2018; Sirota et al., 2003). Yet, direct evidence for such temporally structured hippocampal–cortical interactions at the level of neural populations remains elusive in humans. The simultaneous recording of iEEG and neuronal population activity across multiple MTL regions, together with EEG, enabled us to investigate reciprocal MTL–cortical communication across different spatial and temporal scales.

In a first step we examined the temporal relationship between EEG-detected SOs (n = 34,974; mean ± SEM: 1665.43 ± 101.54), spindles (n = 40,688; mean ± SEM: 1937.52 ± 155.06), and hippocampal SWRs detected in iEEG recordings from AH (for SO and spindle-related ERPs see Fig. S4). In line with previous reports from rodents (Sirota et al., 2003) and humans (Staresina et al., 2023), both spindles and hippocampal SWRs preferentially nested within the up-states of SOs, with SWR density peaking during the first SO up-state (Fig. 3A; spindles: *p* = 0.003, -0.75 to -0.25 s and *p* = 0.002, 0.45 to 0.9 s relative to SO-trough); SWRs: *p* = 0.001, -1.25 to -0.5 s). Moreover, SWRs were significantly coupled to spindle troughs (Fig. S5*, p* = 0.017, –0.1 to 0.15 s; *p* = 0.002, 1.15 to 1.5 s; two-sided paired-samples t-test, corrected across time). To further examine the hierarchical coordination of SOs, spindles, and SWRs, we identified SO–spindle complexes (i.e., events where spindles emerged ±1.5 s around SO down-states). Coupled spindles exhibited consistent phase-locking to SO up-states (Rayleigh test: *z* = 12.96, *p* = 2.0 × 10^−7^; circular mean ± SEM: 45.13 ± 0.15°). Also, SWRs occurring during SO–spindle events (i.e., SO–spindle–SWR triplets) were significantly phase-locked to SO up-states (Rayleigh test: *z* = 4.47, *p* = 9.9 × 10; circular mean = 52.52° ± 0.27). The phase distributions of SO-coupled spindles and SO-coupled SWRs did not differ significantly (Watson–Williams test: *F*(1,40) = 0.16, *p* = 0.693; for details see supplementary text S1; for corresponding dynamics in time-frequency space see Fig. 3 C-E top panels).

**Figure 3.**
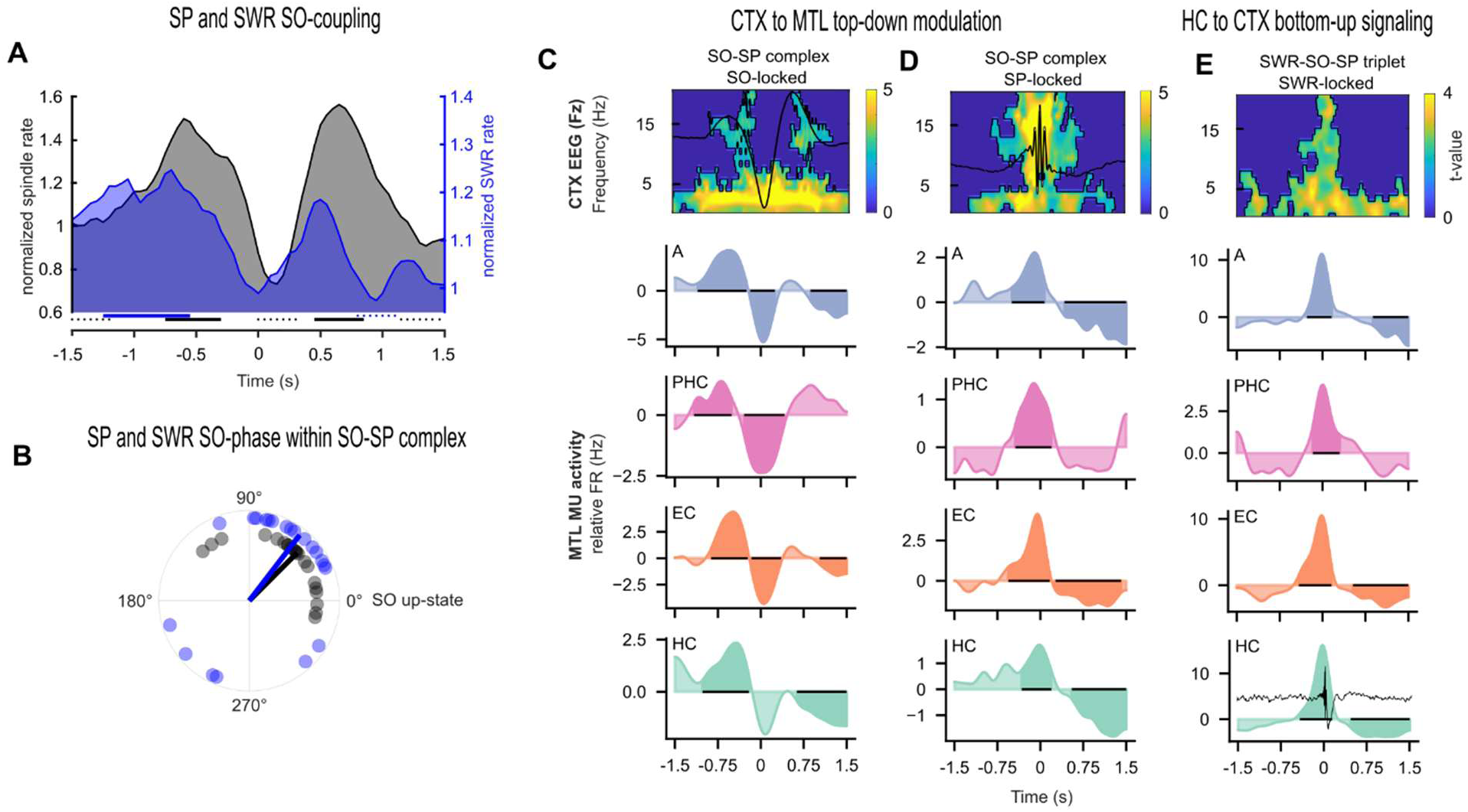
Coupled sleep oscillations shape bidirectional cortical–hippocampal communication. (**A**) Rates of cortical spindles (SP, gray), and hippocampal SWRs (blue) aligned to cortical SO troughs reveals peak occurrence during SO up-states. Spindles exhibited significant clustering during both the first up-state (two-sided dependent-sample t-test; *p* = 0.003, corrected across time) and second up-state (*p* = 0.002), while SWRs peaked during the first up-state (*p* = 0.001); solid lines indicate time points of significantly elevated event rates; dashed lines indicate significantly reduced rates. (**B**) Preferred SO-phases of spindles (blue) and SWRs (red) within SO–spindle complexes. Circular statistics (Rayleigh test) revealed significant nesting of spindles (*z* = 12.96, *p* = 2.0 × 10^-7^) and SWRs (*z* = 4.47, *p* = 9.9 × 10^-3^) in SO up-states. Distributions of preferred phases did not differ (Watson–Williams test: *p* = 0.693), with closely aligned circular means (SPs: 45.13° ± 0.15 and SWRs: 52.52° ± 0.27). (**C**) *Top*: Time–frequency representation (TFR) including ERP of SO–spindle complexes aligned to SO troughs, showing significant increases (*p* < 0.01; unmasked time-frequency bins) of normalized power in the spindle range during the SO upstate (all TFRs: two-sided paired-samples t-test, corrected across time and frequency; see supplementary text S1 for detailed statistics). *Bottom*: PETHs of multi-unit (MU) activity across the MTL reveal top-down entrainment by cortical SOs, inducing states of high excitability during the SO-upstate and suppression during the down-states (shaded areas indicate *p* < 0.05, two-tailed cluster-based permutation test, corrected across time; see Table S2 for detailed statistics). (**D**) *Top*: TFR including ERP of SO–spindle complexes aligned to spindle troughs (*p* < 0.01). *Bottom*: MU activity peaks to spindle troughs across all MTL regions demonstrating global impact of cortical spindles. **(E**) *Top*: TFR of cortical EEG (Fz) aligned to hippocampal SWRs (see ERP in bottom plot) shows significant (*p* < 0.05) power increases in the spindle and SO range, reflecting precise cross-structural coupling. *Bottom*: Strong SWR-linked hippocampal output bursts across the MTL are precisely aligned to cortical SO–spindle complexes.

Having established robust SO–spindle–SWR coordination, closely matching prior findings of hippocampal–cortical interactions (Helfrich et al., 2019), we next asked whether SO–spindle complexes directly modulate MTL population activity. Such modulation would indicate a top-down cortical influence on MTL neural network dynamics. To this end, we aligned multi-unit activity (MUA) from all MTL microwires to an ±1.5 s window centered on the down-state of spindle-coupled SOs. Consistent with rodent studies, spindle-locked cortical SOs induced alternating states of excitability in the MTL, with elevated firing during SO up-states and generalized neuronal hyperpolarization during SO down-states (Fig. 3C; see Table S3 for detailed statistics). Notably, aligning MTL activity to SO-locked spindles revealed peak firing precisely tuned to spindle troughs (Fig. 3D, see Table S3 for detailed statistics), suggesting that cortical SO-spindles exert global, top-down modulation of MTL population dynamics (see Fig. S6 for results in relation to all scalp detected SOs and spindles (i.e., irrespective of SO-spindle events)). Finally, we asked whether hippocampal output would be precisely timed to cortical SO–spindle complexes. We identified SWRs that occurred within ±1.5 s of the SO down-state in SO–spindle complexes and aligned MTL population firing to the SWR centers. SWR events nested within SO–spindle complexes were accompanied by strong population bursts across the MTL, reflecting temporally precise hippocampal output synchronized with cortical network activity (Fig. 3E). Together, these findings support a model of reciprocal cortical–hippocampal communication, enabled by the hierarchical coupling of SOs, spindles, and SWRs across temporal and anatomical scales. This, in turn, raises the question of the functional significance of these synchronized hippocampal–cortical dynamics.

### Coupled hippocampal–cortical sleep oscillations facilitate SWR-locked interregional reactivation of experience-related neurons

As outlined above, contemporary models of systems consolidation propose that the triple coupling of NREM sleep-related oscillations (i.e., SWR-SO-spindle events) might play a key role in memory consolidation (Klinzing et al., 2019; Maingret et al., 2016; Staresina, 2024; Todorova & Zugaro, 2020). The prevailing view is that their coordinated interplay supports both the selective reactivation of recent experiences and the transfer of reactivated memory content from the hippocampus to cortical target regions. However, direct evidence for this mechanism, particularly in humans, remains elusive (Navarrete et al., 2020; Staresina, 2024). Concept cells in the human MTL closely track human behavior and are thought to serve as building blocks of declarative memory (Mackay et al., 2024; Quiroga, 2012). Examining concept neurons tied to recent experience may thus provide a unique window into memory-related processing during sleep. To this end, we identified 248 (mean ± SEM: 11.81 ± 1.32) selective neurons that consistently responded to semantic stimuli across the entire experimental sessions preceding sleep (Niediek et al., 2026; for details see Methods). To investigate the functional role of sleep oscillations and their interplay (in form of SO-spindle-SWR triplets) for memory reactivation, we first hypothesized that hippocampal SWRs would promote the selective reactivation of memory traces from recent experiences (see also Niediek et al., 2026), reflected in the coordinated co-firing of semantically tuned neurons. This selective ensemble reactivation could serve to differentiate relevant from irrelevant information, offering a potential mechanism by which SWRs prioritize behaviorally significant information for cortical transfer within the sequential coupling.

First, we compared the co-activation strength of task-related selective neuron pairs to that of non-responsive neurons (n = 849; mean ± SEM: 40.33 ± 5.23) during SWRs (± 50ms). To isolate SWR-specific effects on neural co-firing, while accounting for firing rate variability and stereotypic intrinsic firing, we normalized the cross-correlograms (CCGs) by subtracting SWR-triggered shift predictors and a baseline CCG (also shift-predictor corrected (Staresina et al., 2023, see Methods). Only neuron pairs whose co-activation levels exceeded the baseline co-firing after correction were included in the final analyses. Given the widespread hippocampal-to-cortical information flow during SWRs (Fig. 2D and 3E), we assessed SWR-locked co-activation strength by region, analyzing neuron pairs both within the hippocampus and in downstream MTL regions. As expected, task-related neurons in the AH showed significantly stronger co-activation than non-responsive neurons (AH: *p* = 6.6 × 10^−4^; Wilcoxon rank-sum test, FDR-corrected; Fig. 4B), extending prior observations on hippocampal neural reactivation (Niediek et al., 2026) with metrics that specifically resolve SWR-driven coactivation gains, while no significant difference was found in the posterior hippocampus (*p* = 0.18). However, the selective co-reactivation of task-related neuron activity extended beyond the hippocampus, with robust effects observed in the A (*p* = 1.8 × 10^−3^) and PHC (*p* = 5.6 × 10^−20^; note that the EC was included in Fig. 4B for illustrative purposes only, as it did not exhibit enough co-activation pairs and was therefore excluded from the statistical analysis). These results suggest that SWRs drive the selective reactivation of experience related information across both hippocampal and downstream cortical regions.

**Figure 4.**
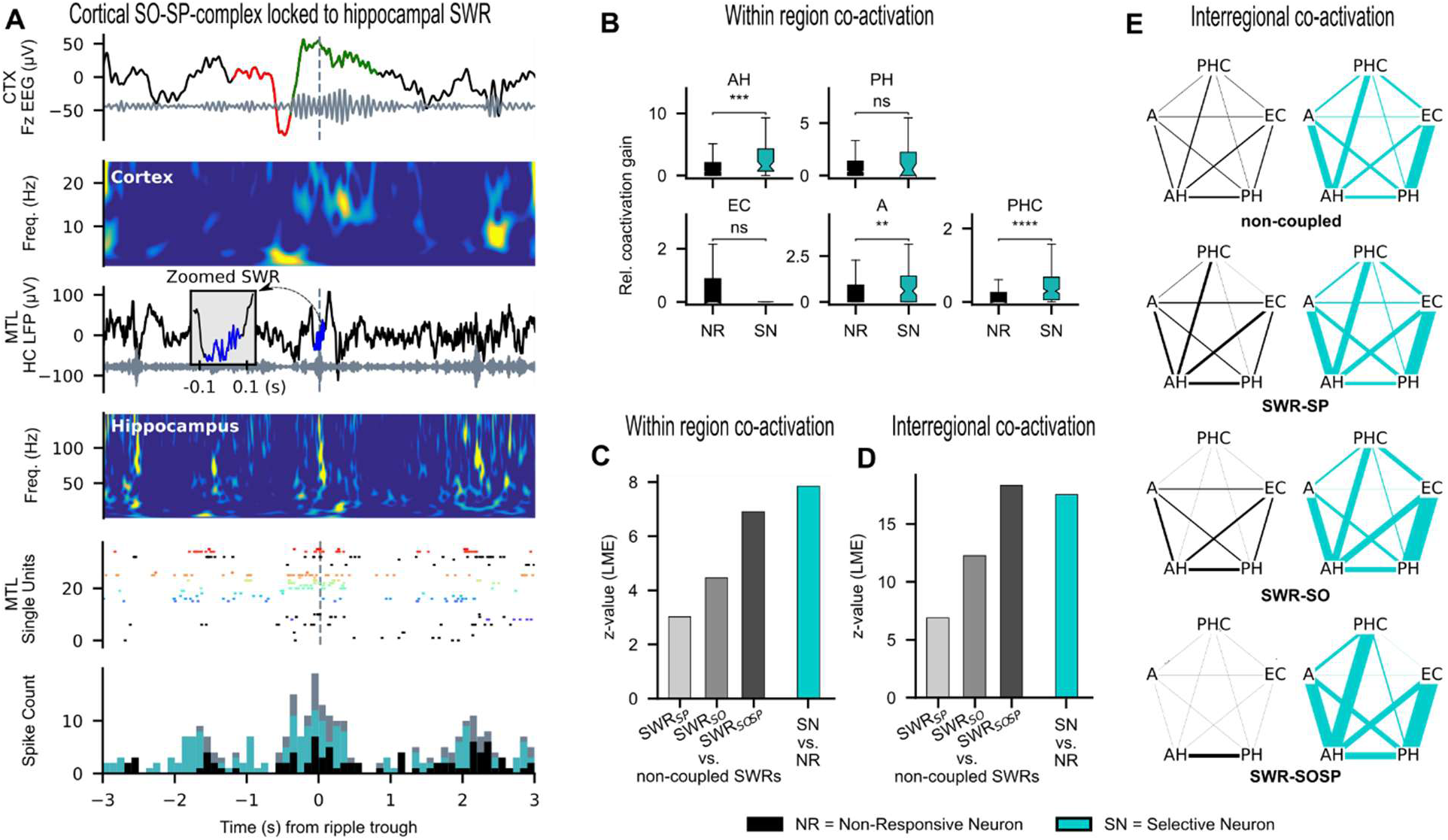
Selective-neuron coactivation is modulated by hippocampal–cortical coupling. (**A**), Example of a single cortical SO–spindle complex coupled to a hippocampal SWR (time 0 s, dashed vertical lines). Top: EEG shows a slow oscillation (red) and spindle (green), with corresponding time–frequency representation (0.1–25 Hz). Middle: SWR event in hippocampal iEEG, with corresponding time–frequency representation (0.1–150 Hz). Bottom: Raster plot of MTL neurons, with selective neurons shown in color and non-responsive neurons in black. Below, the population spike histogram shows firing rates of selective neurons (cyan) and non-responsive neurons (black), with the cumulative population activity overlaid in grey. Selective neurons show increased firing during SWR events(0s) SWR (0 s). (**B**), SWR-triggered coactivation gain for intra-regional neuron pairs. experience related (selective) neuron pairs (cyan) exhibit significantly stronger coactivation than non-responsive pairs (black) in AH (see related results in Niediek et al. 2026), and in downstream regions (A and PHC). (**C**), Bars show fixed-effect z-values from a linear mixed-effects model, fitting coactivation gain across coupling conditions (left, grey bars) and neuron types (right, cyan bar). SWR coupling effects are expressed relative to non-coupled ripples and averaged across cell types, whereas the cell-type effect is averaged across SWR coupling conditions. Coupling to cortical events significantly enhances coactivation, with the strongest gain during SWR-SO-spindle triplets. Selective neurons show significantly higher coactivation across all coupling states. (**D**), Same analysis as in C), now for cross-regional neuron pairs (e.g., AH–PHC), confirming that SWR–cortical coupling enhances selective coactivation of experience-related neurons across regions. (**E**), Schematic illustration of cross-regional coactivation strength (edge thickness) for selective (cyan) and non-responsive (black) neuron pairs across coupling conditions. Across conditions, from uncoupled SWRs (top) to SWR-SO-spindle triplets (bottom), coactivation among non-responsive neurons decreases, while coactivation among experience related (selective) neurons becomes increasingly pronounced, suggesting selective systems-level reactivation.

Given the assumed role of cortical SO-spindle complexes in memory consolidation (Todorova & Zugaro, 2020) and their active role in shaping MTL firing dynamics (Fig. 3) (Sirota et al., 2003), we next asked whether coincidence of these oscillations would facilitate neuronal co-activation. To test this systematically, we compared co-activation strength during uncoupled SWRs with SWRs that were coupled to cortical events, namely spindles (SWR-SP), SOs (SWR-SO), or full SO–spindle complexes (SWR-SO-SP).

To examine the influence of neuronal response type and event coupling on co-activation strength, we employed a linear mixed-effects model (random intercepts for patients, estimated using maximum likelihood). Analyses were restricted to within-region neuron pairs. Neurons were categorized as selective or non-responsive (model baseline), and coupling conditions were defined as uncoupled (model baseline), SWRs coupled to spindles (SWR-SP), SWRs coupled to SOs (SWR-SO), or SWR-SO-SP triplets.

Strikingly, hippocampal coupling to cortical oscillations was associated with robust increases in co-activation strength relative to uncoupled SWRs. This effect was strongest for SWRs coupled to SO-spindle complexes (ß = 0.35, SE = 0.05, z = 6.91, p < 0.001), which elicited the highest co-activation across all conditions. Significant increases were also observed for coupling with solitary SOs (SWR-SO: ß = 0.23, SE = 0.05, z = 4.46, p < 0.001) and solitary spindles (SWR-SP: ß = 0.15, SE = 0.05, z = 3.02, p = 0.003), reflecting a graded enhancement in line with the level of cortical engagement (see Fig. 3C).

Importantly, selective neurons exhibited a pronounced increase in co-activation strength relative to non-responsive units across all coupling conditions (ß = 0.52, SE = 0.07, z = 7.85, p < 0.001). No interaction terms were included in the final model, as interaction effects were not significant in preliminary analyses, indicating that the enhanced co-activation associated with neuronal selectivity was preserved across all oscillatory coupling conditions.

While the above analyses examined co-activation within individual hippocampal or adjacent regions, we finally explored whether neuronal ensembles also exhibit coordinated reactivation across distributed MTL circuits. To this end, we extended our co-activation analysis to cross-regional neuron pairs (e.g., AH-PHC) using the same linear mixed-effects modeling framework (random intercepts for patients, maximum-likelihood estimation). Remarkably, selective neurons again exhibited significantly stronger cross-regional co-activation than non-responsive units across coupling conditions (ß = 0.44, SE = 0.03, z = 17.54, p < 0.001).

Co-activation strength was significantly elevated across all coupling conditions relative to uncoupled events, with the strongest effect observed during SWR-SO-spindle triplets (ß = 0.31, SE = 0.02, z = 18.32, p < 0.001), followed by SWRs coupled to solitary SOs (ß = 0.20, SE = 0.02, z = 12.26, p < 0.001) and solitary spindles (ß = 0.12, SE = 0.02, z = 6.88, p < 0.001). No interaction terms were included in the final model, as interactions between unit type and coupling condition were not significant in preliminary analyses, indicating that the advantage of selective neurons was preserved across all oscillatory coupling conditions.

For a qualitative illustration of the network activity underlying these effects, see Fig. 4E. Together, these findings indicate that experience-related neurons form distributed cross-regional ensembles that are selectively reactivated during SWRs. Coupling to cortical oscillations (particularly SO-spindle complexes) robustly amplifies these reactivation processes, pointing to a synergistic hippocampal-cortical mechanism in support of memory consolidation.

## Discussion

Our findings provide evidence on a single-neuron level that SWRs, in coordination with cortical SOs and spindles, orchestrate the selective reactivation of experience-related neurons across the human medial temporal lobe (MTL). Leveraging rare multi-site recordings during natural sleep, we demonstrate that SWRs not only direct hippocampal-to-cortical information flow but also preserve behaviorally relevant content across distributed MTL structures. Reactivation is strongest when hippocampal SWRs are precisely aligned with cortical SO–spindle complexes, suggesting that the hierarchical coupling of sleep-related oscillations serves as a temporal scaffold for systems-level memory consolidation.

Hippocampal SWRs have been causally linked to memory consolidation in rodents (Fernández-Ruiz et al., 2019; Girardeau et al., 2009; Maingret et al., 2016) and correlate with retrieval-related reactivation during wake in humans (Kunz et al., 2024; Norman et al., 2019; Schreiner et al., 2024; Vaz et al., 2020). Recent models of memory consolidation (Diekelmann & Born, 2010; Klinzing et al., 2019) suggest that SWRs transmit “chunked” mnemonic content across brain regions via multiplexed signaling (Buzsáki, 2015; Logothetis et al., 2012; Swanson et al., 2020; Todorova & Zugaro, 2020). Yet, evidence for such directed, brain-wide information flow has been limited in rodent models (but see (Nitzan et al., 2022) and entirely lacking in humans. Here, we show that SWRs initiate a stepwise propagation of activity along the hippocampal longitudinal axis and towards downstream targets, including the entorhinal and parahippocampal cortices and the amygdala. This anterior-to-posterior flow mirrors some recent results in animal models (e.g., (Patel et al., 2013)), suggesting that functional specialization along the hippocampal axis may guide the direction of information transfer (Nitzan et al., 2022). Our data positions the anterior hippocampus as a key hub for intra- and extra-hippocampal communication, raising the question of whether hippocampal output contains information that can be read-out at downstream regions.

Human concept neurons encode semantic identity independent of context (Bausch et al., 2026; Rey et al., 2025) and are thought to represent the “what” component of episodic memory, complementing the spatial and temporal coding seen in rodent place cells (Mackay et al., 2024; O’Keefe, 1976; O’Keefe et al., 1971). Unlike place cells, concept neurons are distributed across several MTL regions (Quiroga, 2012), raising the question of how such scattered representations are coordinated during sleep. Our data suggest that reactivation of complex memory content requires the large-scale synchronization of semantically tuned neurons, across both hippocampal and downstream structures, timed by cortically driven sleep dynamics. Each MTL subregion may contribute distinct facets to the memory trace, such as affective salience from the amygdala (Girardeau et al., 2017; Mormann et al., 2011, 2015) or contextual and spatial detail from the parahippocampal cortex (Kunz et al., 2024; Mackay et al., 2024), pointing toward a systems-level integration process.

While previous work has mostly focused on single hippocampal–cortical connections (but see (Miyawaki & Mizuseki, 2022)), we found that experience-related neurons exhibit coordinated reactivation patterns across multiple MTL regions during SWRs, including distal sites like the amygdala and parahippocampal cortex. This supports the idea that hippocampal output carries behaviorally relevant content that can be read out and integrated across distributed networks. Importantly, SWRs do not occur in isolation, but are embedded within a broader temporal hierarchy of cortical rhythms (Latchoumane et al., 2017; Maingret et al., 2016; Sirota et al., 2003; Staresina et al., 2015). In particular, the coordinated coupling of SOs, spindles, and SWRs has been causally linked to consolidation efficacy (Maingret et al., 2016). SOs may gate hippocampal output by transiently increasing brain wide cortical excitability and facilitating SWR–spindle coupling (Sirota et al., 2003). In line with this model, we observed that MTL population firing was modulated by SO–spindle cycles, with SO up-states enhancing and down-states suppressing activity. Spindles further timed firing to their troughs (Averkin et al., 2016; Sirota et al., 2003), supporting the idea that cortical oscillations shape MTL dynamics in a top-down manner. Once routed to the MTL, these cortical inputs may engage local circuit mechanisms: it has recently been shown that local SOs, spindles, and SWRs detected in the MTL can sequentially constrain co-firing within progressively narrower time windows, potentially enabling synaptic plasticity (Staresina et al., 2023). Yet, how these interactions coordinate memory-specific content across hippocampal–cortical circuits remained unclear.

Prior studies, including our own, have typically focused on isolated oscillatory components, lacked direct evidence of experience-dependent reactivation (but see Niediek et al., 2026, were limited to the MTL, or used methods without single-neuron resolution (e.g., (Helfrich et al., 2019; Maingret et al., 2016; Peyrache et al., 2009; Schreiner et al., 2024; Schreiner & Staudigl, 2020; Staresina et al., 2023). Here we directly assessed the functional significance of coupled sleep oscillations for information transfer, by examining SWR-related co-activation among concept neurons. Co-activation strength increased significantly when SWRs were coupled with cortical oscillations, with the strongest effects during triple events. This graded enhancement, from spindles to SOs to full SO–spindle events, was consistent across hippocampal and extra-hippocampal regions. Notably, task-related selective neurons consistently showed higher co-activation than non-responsive units, independent of coupling condition. This suggests that SWRs are the primary driver of selectivity, while SO–spindle coordination enhances the effectiveness of reactivation, possibly by aligning it with cortical windows of optimal plasticity. Extending beyond regional analysis, we observed selective co-activation across anatomically distributed neuron pairs, further supporting the idea of SWR-driven, large-scale memory reactivation. These effects were again amplified when all three key oscillations of NREM sleep coincided, reinforcing the view that cortical SO–spindle complexes serve as a temporal scaffold for hippocampal output.

Taken together, these findings support a model of cortically gated hippocampal reactivation, in which SO–spindle complexes provide a temporal scaffold for selectively routing hippocampal memory content to downstream targets during windows of putative heightened cortical receptivity. In this framework, cortical rhythms do not merely accompany hippocampal output, but actively structure it: tagging relevant content, aligning neuronal excitability, and facilitating interregional plasticity. This model integrates bottom-up reactivation with top-down control and provides a mechanistic account of how distributed elements of an experience are selectively consolidated into lasting memory traces.

## Methods

### Participants and Recordings

We analyzed 21 full-night sleep recordings (mean 2.1 ± 0.31 per patient) from 10 individuals (5 female, 5 male, aged 20–62 years, mean age 39.9 years) undergoing intracerebral monitoring for pharmacologically intractable epilepsy. All participants had been implanted with clinical depth electrodes as part of their standard pre-surgical evaluation. These data represent a subset of the data (21 of 40 sessions) reported in Niediek et al. (2026), restricted to patients in whom EEG, clinical iEEG, and single-neuron activity were simultaneously recorded on the same acquisition system, yielding the temporal alignment necessary for the present analyses. The study was approved by the Medical Institutional Review Board of the University of Bonn, and all participants provided written informed consent prior to inclusion. Multisite iEEG and single-neuron recordings were obtained using Behnke-Fried hybrid macro–micro depth electrodes (AdTech, Racine, WI). The clinical implantation followed a bitemporal scheme, with two (6 patients) to three (4 patients) depth electrodes targeting each hippocampus, and one electrode each placed in the entorhinal cortex, parahippocampal cortex, and amygdala of each hemisphere. To enable multi- and single-unit recordings, each depth electrode was equipped with a bundle of nine microwires (including one reference wire), inserted through the hollow shaft and extending approximately 4 mm beyond the tip of the clinical depth electrode. Additionally, scalp EEG (F3, F4, Fz, C3, C4, O1, O2, Cb1, Cb2), electrooculography (EOG), and submental electromyography (EMG) were recorded in parallel for polysomnography. Sleep scoring was performed according to AASM criteria (Iber et al., 2007). All signals were recorded and amplified using a 256-channel Neuralynx ATLAS system (Bozeman, MT). Scalp EEG and intracranial EEG (iEEG) were band-pass filtered (0.1–600 Hz) and sampled at 2,048 Hz. Microwire signals were band-pass filtered (0.1–9,000 Hz) and sampled at 32 kHz. All data were digitally stored for offline analysis.

### Study Design

The study design is described in detail in (Niediek et al., 2026). In brief, a typical session consisted of two subsequent testing days. In the morning of Day 1, participants first underwent a stimulus-screening session to identify visually responsive neurons and their preferred image stimuli. In the evening of Day 1, participants completed an episodic memory task involving the learning and immediate recall of a narrative (“Fotonovela”) involving the previously identified stimuli. To assess the recording stability and response selectivity of the identified neurons, brief screening sessions were performed before and after the memory task in the evening of Day 1 and again following delayed recall after sleep in the morning of Day 2 (see also Fig. 1 B for experimental procedure).

### Episodic Memory Task: “Fotonovela”

The “Fotonovela” task was designed to induce task-related, behaviorally linked activity of concept neurons through a narrative/story built around response eliciting images identified in the morning screening (Niediek et al., 2026). The participant-tailored story was presented as a sequence of 8 to 10 slides (mean ± SEM: 9.2 ± 0.17), with each slide featuring a response-eliciting picture with its corresponding written name and 2–4 connecting sentences. These connecting sentences were used to establish an episodic link between the concepts of the previous, current, and next slide. All identified response-eliciting images were included in the task. Participants viewed the slides in a self-paced manner and were instructed to memorize the order of the concept images. Afterward, they recalled the story by naming the concepts in sequence, with feedback provided after each correct response. Recall accuracy was scored based on the correct naming of concepts in order. Patients were encouraged to recall the linking sentences, but only correct concept naming was required for a correct score. Participants repeated the recall task until they successfully recalled the entire story six times. In the morning of Day 2, they completed a final recall session.

Furthermore, a mini-screening was run to confirm neuronal responses to the preferred pictures. Additionally, written names describing each of these pictures were presented in isolation to characterize invariance of responses. This mini-screening was run 3 times: In the evening of Day 1 both before and after the Fotonovela task, and in the morning of Day 2 after the final recall session.

### Selective neurons during a full night of sleep

As detailed in (Niediek et al., 2026) visually responsive neurons and their corresponding eliciting stimuli were identified on the morning of Day 1 using previously described procedures (Quiroga et al., 2005). In brief, 100–150 images, including persons (celebrities, patient’s friends, and relatives), animals, scenes, and objects, were each presented to the patient six times. Subsequently, up to ten response-eliciting images for the memory task were selected. To assess semantic invariance, additional screening sessions were conducted using only the selected images and their corresponding written labels. A unit was considered responsive to a stimulus based on a combination of a selectivity index and a response score.

The selectivity index *SI_i_* was quantified using a modified z-score, reflecting how selectively a neuron responds to stimulus *i* relative to all other stimuli. It is computed as the difference between the neuron’s mean spike count for stimulus *i* and the average spike count across all other stimuli, normalized by the standard deviation of the responses to the other stimuli: 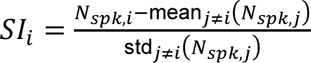

Here, *N_spk,i_* denotes the average spike count across all presentations of stimulus *i*, while the mean and standard deviation are calculated over all other stimuli (*j ≠ i*). A higher SI indicates stronger selectivity for that stimulus.

Response score RS_i_ was assessed following (Mormann et al., 2008) by comparing spike counts during the 0–1000 ms post-stimulus interval to a 500 ms pre-stimulus baseline, using Wilcoxon signed-rank tests in 19 overlapping 100 ms bins. The minimum p-value across bins, corrected for multiple comparisons using the Benjamini–Hochberg method, defined the response score for each stimulus. A stimulus-unit pair was classified as a response if it met at least one of the following criteria (Niediek et al., 2026):

(1) SI_i_ ≥ 5 and RS_i_ < 1;
(2) SI_i_ ≥ 3 and RS_i_ < 0.5;
(3) SI_i_ ≥ 1 and RS_i_ < 0.001.

Units responding to a stimulus in either the evening or morning or both were labeled responsive units. Neurons that showed responses to multiple images were removed from subsequent analyses, and only those neurons that responded to a single image within the mini-screening set were classified as selective neurons. Responses were deemed invariant if a unit responded to both the image and the corresponding written label of a single concept, or to the image alone with a selectivity index ≥ 3 for the corresponding label. Out of 248 selective neurons, 71 (28,6%) neurons showed semantic invariance.

### Neuronal recordings during human sleep

Neuronal spikes were extracted from individual microwires using Combinato, a Pyhton-based spike sorting framework optimized for long-term single-neuron recordings under noisy conditions(Niediek et al., 2016). For spike detection, signals were first band-pass filtered (300–3,000 Hz) and then thresholded. To ensure reliable identification of neural activity, a multi-step artifact rejection was applied using Combinato’s default settings. First, 500-ms windows (with 250-ms overlap) containing more than 100 detected events (i.e., firing rate > 200 Hz) were excluded to eliminate noise-induced bursts. Second, events exceeding 1 mV in amplitude were removed as non-physiological. Finally, 3-ms bins (1.5-ms overlap) showing activity on more than 50% of channels were excluded, to reduce artificial co-activations from global artifacts.

To identify single units among this artifact-cleaned multi-unit activity, spike sorting was carried out using Combinato with default settings, except for adjusting the parameters to Smin = 25 and Cstop = 1.6. Manual refinement of the clustering was limited to channels showing stimulus-responsive activity, identified during the morning screening. For each detected unit, raster plots and PETHS were generated and reviewed to assess visual responsiveness. In those responsive channels, further curation was performed using Combinato’s graphical interface—this included discarding residual artifacts and merging clusters that appeared to represent the same neuron (overclustering). Over the course of an overnight recording—often exceeding 12 hours—neuronal signals can be intermittently lost due to microwire drift, tissue movement, or transient noise. To ensure that only stably recorded neurons were included in the analysis, we assessed unit stability by dividing the recording into 5-minute segments and calculating the proportion of segments in which the firing rate exceeded 5% of the unit’s mean firing rate across the entire night. Units with a recording stability below 90% were excluded from further analysis.

### Scalp EEG (sEEG) and iEEG analyis: Artifact rejection and event detection

Scalp EEG and iEEG data were analyzed using MATLAB (R2024a) and FieldTrip (version 20231015, (Oostenveld et al., 2011)). To detect artifacts (e.g. sharp transients) and interictal epileptic discharges (IEDs), an established algorithm (Staresina et al., 2015) was implemented. Two auxiliary transformations of the raw signal were derived: one accentuating high-frequency components via a 250-Hz high-pass filter, and another capturing transient fluctuations through the first temporal derivative of the raw signal. These, along with the raw signal, were independently z-scored within each sleep stage to normalize for stage-specific variance. For artifact detection, a dual-threshold criterion was employed: data points exceeding a z-score of 6 in any of the three signals were flagged, as were those in which the raw signal, together with either the band-passed or derivative signal, concurrently exceeded a z-score of 4. Identified artifacts were subsequently extended by 0.5 seconds on either side to account for potential boundary effects and excluded from all further analyses. Cortical SOs and spindles, as well as hippocampal SWRs were detected using established procedures applied to artifact-free NREM epochs.

SOs were identified by first band-pass filtering the EEG from Fz between 0.3 and 1.25 Hz using a two-pass finite impulse response (FIR) filter, with the filter order set to three cycles of the lower cutoff frequency. Then, all zero-crossings were identified in the band-pass filtered signal, and candidate SO events were defined as segments spanning two consecutive positive-to-negative zero-crossings - corresponding to a full cycle from downstate to upstate. Only events with durations between 0.8 and 2 seconds were kept for further analysis. Finally, the amplitude of each remaining SO candidate was quantified by measuring both the trough depth and the trough-to-peak difference. Events were classified as SOs if both measures exceeded the mean plus 1.5 SD across all candidates.

To detect spindles, the signal from Fz (C4 in one patient due to noise) was band-pass filtered between 12 and 16 Hz (two-pass FIR band-pass filter, order = three cycles of the low-frequency cutoff), after which the root mean square (RMS) envelope was computed using a 200-ms sliding window and subsequently smoothed with sliding window of the same size. Candidate spindle events were defined as segments in which the smoothed RMS signal exceeded a threshold set at 1.4 SD above the mean RMS value across all NREM epochs, lasting between 0.4 and 3 seconds. Events of interest exceeding an upper threshold of 9 SD were excluded to avoid contamination by high-amplitude transients. In addition, only events containing at least six oscillatory cycles in the raw EEG signal were kept for further analysis. Finally, to exclude false positives, candidate spindles were required to exhibit a prominent peak in the TFR between 10 to 18 Hz within a window of 0.8 seconds around the spindle center.

SWR detection then followed a parallel approach to that used for spindles, with adaptations for activity in higher frequencies. For SWR detection, iEEG signals of the deepest and second-deepest macro contacts on electrodes targeting the hippocampus were referenced as a bipolar montage. For computational efficiency iEEG data were first downsampled to 1024 Hz and then band-pass filtered between 80 and 120 Hz. Subsequently, the RMS envelope was computed using 20-ms sliding windows, followed by smoothing with the same window length. Candidate events were identified when the smoothed RMS signal exceeded a threshold defined as the mean plus 3 SD of the RMS across all periods within each NREM stage. Candidate events exceeding an upper threshold of mean plus 9 SD were excluded to avoid contamination from high-amplitude transients. Only events lasting between 38 ms (equivalent to three cycles at 80 Hz) and 500 ms were considered. To further ensure oscillatory integrity, each SWR was required to contain at least three algorithmically detected cycles in the raw signal. Finally, to reduce false positives, candidate SWRs were required to exhibit a clear spectral peak in the time–frequency representation between 75 and 125 Hz within a ±0.25-s window centered on the SWR midpoint.

### Coupling analyses

Peri-event time histograms (PETHs) were computed for all combinations of sleep oscillations within a ±1.5-s window centered on the trigger event. Event times were defined as the SO downstate (negative peak), spindle trough, or SWR peak amplitude for all analyses. PETHs were constructed using 50-ms bins, normalized by event number, and smoothed by convolving with a 100-ms Gaussian kernel. Coupled events, such as SO-Spindle complexes, were defined by event hits within the ±1.5-second time window of the PETH.

PETHs relating sleep-oscillation timing to multi-unit activity (MUA) followed the same procedure, except using 1-ms bins to resolve the finer temporal resolution of spike timing. For further event or regionwise analysis, PETHs were averaged across hemispheres. Statistical significance was assessed using a nonparametric, two-sided cluster-based permutation test based on paired-samples *t* statistics (1,000 permutations, *P* < 0.05), with correction for multiple comparisons over time (Maris & Oostenveld, 2007), as implemented in MNE-Python (Gramfort et al., 2013), where the observed PETH was compared to a random distribution generated by 1,000 random shuffles of the PETH.

To directly assess event co-occurrences based on spectral power, we performed time-frequency analysis using FieldTrip (Oostenveld et al., 2011) functions on signal epochs spanning ±3 sec around event centers. Time–frequency analysis was performed using a sliding-window approach, with window lengths set to five cycles of each frequency. Frequencies ranged from 0.5 to 20.5 Hz in 1-Hz steps for cortical events, and the range was extended up to 200.5 Hz for hippocampal SWR events. Each data segment was tapered by pointwise multiplication with a Hanning window prior to Fourier transformation to reduce spectral leakage. Power estimates were then z-scored to a pre-event baseline (−3 to –1.5 s). Extended time windows were used to improve resolution of low-frequency components within the analysis window of interest (−1.5 to 1.5 s) and to mitigate edge artifacts.

Finally, we examined the phase relationships between cortical SOs, spindles and hippocampal SWR events. For event timings relative to the SO phase, cortical EEG data were filtered between 0.5 and 2 Hz using a two-pass Butterworth bandpass filter, and the analytical signal was then obtained via Hilbert transform. The instantaneous phase angle of the cortical signal was then extracted at the time of each cortical spindle or hippocampal SWR event. The preferred phase of the SO-to-event coupling was computed by calculating the circular mean of the individual event phases using the Circular Statistics toolbox for Matlab (Berens, 2009). A Rayleigh test was applied to assess whether the distribution of preferred phases exhibits significant concentration around a particular phase, indicating a preferred phase for the events. To specifically test for event nesting within SO up-states, a V-test against phase 0 (corresponding to the SO upstate) was used. Finally, phase coupling between spindles and SWRs with respect to the SO phase was assessed using a circular paired samples t-test. A non-significant result indicates substantial overlap between their preferred phases relative to SOs.

### SWR-triggered information flow

To systematically investigate SWR-associated information flow along the hippocampal longitudinal axis as well as the hippocampal-to-cortical axis, time lags of peak firing during hippocampal SWRs in different downstream regions were assessed. SWR-triggered spike-count histograms for single/ multi units were computed in 1-ms bins within a ±0.5-s window centered on the hippocampal SWR peak (maximum amplitude), smoothed with a 3-ms Gaussian kernel, and normalized by subtracting the mean firing rate during a pre-event histogram of the same length, centered 1 s before the event. SWR-associated burst timings were defined as peak firing within a ±75-ms window around the SWR center (mean SWR duration: 56.9 ± 0.08 ms). Time-lag differences in burst-timing distributions between seed (e.g., anterior hippocampus) and target regions were then statistically assessed using nonparametric statistics.

### Relative Coactivation Gain

To assess SWR-associated co-activation, we computed cross-correlograms (CCGs) for all pairwise combinations of single/ multi units within the same behavioral category (e.g., selective neurons, non-responsive neurons), both within and across regions (e.g., AH–AH, AH–PHC).CCGs were calculated in ±50-ms windows around SWR peaks using 1-ms bins and smoothed with a 10-ms Gaussian kernel. To isolate SWR-specific coactivation, multiple correction steps were applied. First, a SWR-centered CCG was corrected using a shift predictor (a copy of the CCG now constructed from n+1 trial-wise shuffling of spike trains) to account for systematic coincident firing arising from neurons with similar rate profiles during SWRs, independent of precise pairwise spike timing. Second, a baseline CCG was computed from a window centered 1 s prior to the SWR peak and subsequently shift-predictor-corrected. Third, the final SWR-evoked CCG was obtained by subtracting the corrected baseline from the corrected SWR-centered CCG, isolating co-activation specifically time-locked to SWRs while reducing covariation driven by baseline firing rates. Final CCGs were then scaled to counts per second and the peak co-firing in the CCG was extracted as final measure of relative co-activation gain. For statistical analysis, we used FDR-corrected Wilcoxon rank-sum tests to compare co-activation strengths between non-responsive and selective neuron pairs within each region (Fig. 4B). To assess the effect of SWR coupling with scalp events, we applied a linear mixed-effects model with fixed factors: coupling type (SWR-SP, SWR-SO, SWR-SO-SP; non-coupled SWRs as baseline) and neuron type (selective vs. non-responsive; with the latter as baseline), and included patient as a random intercept to account for inter-subject variability.

