## Supplemental Material for "Sequential coupling of sleep oscillations enables concept-neuron reactivation and supports information flow across the human hippocampal-cortical circuit"

### SUPPLEMENTARY INFORMATION

| Stage | Mean Percentage (%) | Mean Duration (h) | SEM Duration (h) |
|---|---|---|---|
| **N1** | 23.07 | 1.59 | 0.12 |
| **N2** | 41.09 | 2.95 | 0.22 |
| **N3** | 18.76 | 1.26 | 0.08 |
| **REM** | 17.09 | 1.25 | 0.12 |
| **Total Sleep Time** | 100 | 7.06 | 0.34 |

**Table S1.** Overview of sleep architecture including NREM (N1, N2, N3) and REM epochs.

| Event | Total | Mean | SEM |
|---|---|---|---|
| **SO** | 34974 | 1665.43 | 101.54 |
| **SP** | 40688 | 1937.52 | 155.06 |
| **SO_SP** | 10514 | 500.67 | 46.2 |
| **SWR** | 50118 | 1261.19 | 136.08 |
| **SWR_SP** | 13902 | 331 | 29.85 |
| **SWR_SO** | 11292 | 268.86 | 22.86 |
| **SWR_SOSP** | 4620 | 110 | 12.4 |

**Table S2.** Overview of detected events. Mean and SEM per session.

| Event/Region | Time Window (ms) | p |
|---|---|---|
| SO-SP Complex (SO-trough locked) | | |
| **HC** | -939 to -214 | p = .014 |
| **HC** | 645 to 1499 | p = .001 |
| **EC** | -868 to -233 | p = .001 |
| **EC** | -168 to 365 | p = .001 |
| **EC** | 1042 to 1499 | p = .040 |
| **PHC** | -1096 to -460 | p = .015 |
| **PHC** | -294 to 426 | p = .013 |
| **A** | -1105 to -250 | p = .001 |
| **A** | -189 to 255 | p = .028 |
| **A** | 882 to 1499 | p = .001 |
| SO-SP Complex (SP-trough locked) | | |
| **HC** | -325 to 188 | p = .005 |
| **HC** | 581 to 1499 | p = .001 |
| **EC** | -481 to 159 | p = .003 |
| **EC** | 238 to 1360 | p = .001 |
| **PHC** | -490 to 215 | p = .003 |
| **A** | -534 to 109 | p = .003 |
| **A** | 417 to 1499 | p = .001 |
| *SO-SP Complex (SWR-peak locked)* | | |
| **HC** | -397 to 150 | p = .009 |
| **HC** | 462 to 1499 | p = .001 |
| **EC** | -409 to 160 | p = .002 |
| **EC** | 528 to 1499 | p = .001 |
| **PHC** | -166 to 265 | p = .033 |
| **A** | -291 to 147 | p = .002 |
| **A** | 680 to 1499 | p = .001 |
| *All SOs (SO trough locked)* | | |
| **HC** | -963 to -303 | p = .001 |
| **HC** | 962 to 1499 | p = .001 |
| **EC** | -883 to -244 | p = .001 |
| **EC** | -202 to 357 | p = .001 |
| **EC** | 399 to 982 | p = .007 |
| **EC** | 1156 to 1499 | p = .015 |
| **PHC** | -1114 to -421 | p = .003 |
| **PHC** | -302 to 400 | p = .001 |
| **PHC** | 522 to 1499 | p = .008 |
| **A** | -1018 to -279 | p = .001 |
| **A** | -223 to 248 | p = .001 |
| **A** | 373 to 698 | p = .006 |
| **A** | 946 to 1499 | p = .001 |
| *All SPs (SP trough locked)* | | |

| | | |
|---|---|---|
| **HC** | -284 to 200 | p = .034 |
| **HC** | 330 to 1499 | p = .001 |
| **EC** | 239 to 1245 | p = .001 |
| **PHC** | -642 to 87 | p = .002 |
| **A** | -311 to 108 | p = .025 |

**Table S3.** Two-sided paired-samples t test, corrected over time, for MTL multi-unit FR modulation analysis (main text Fig.3 C-E) locked to different events within SO-SP complexes and for all SO and SP events (see Fig.S6 A and B).

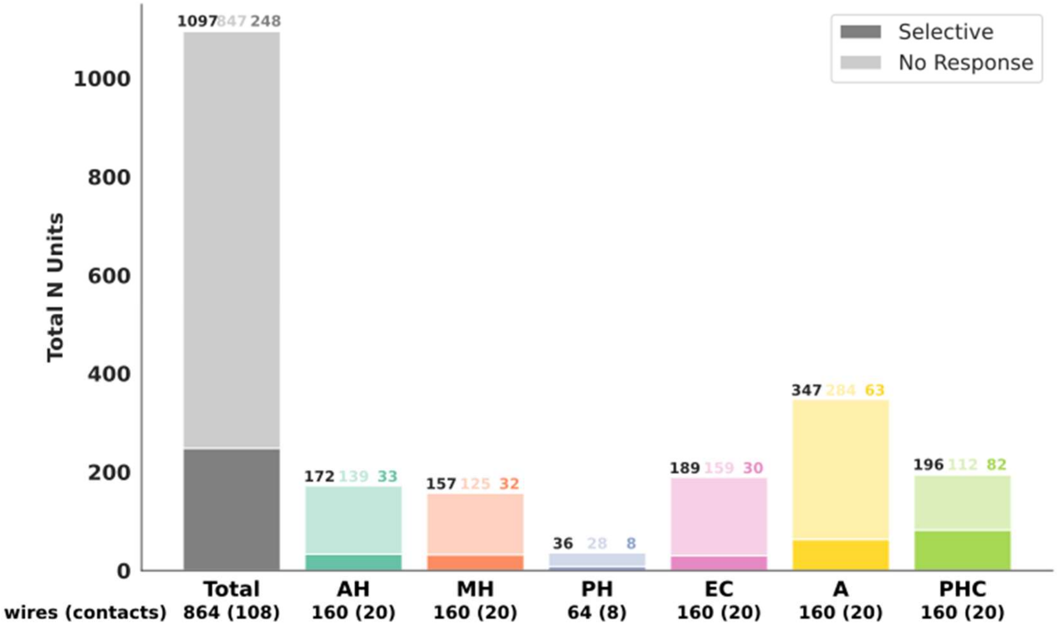


**Figure S1.** Overview of numbers of microwire and bipolar channels, numbers of recorded non-responsive neurons (faded) and recorded selective neurons (solid) in total and per region. Regions are labeled according to the physicians' anatomical assignments based on visual inspection following implantation. Middle hippocampus (MH) and posterior hippocampus (PH) were combined and are referred to throughout the study as PH.

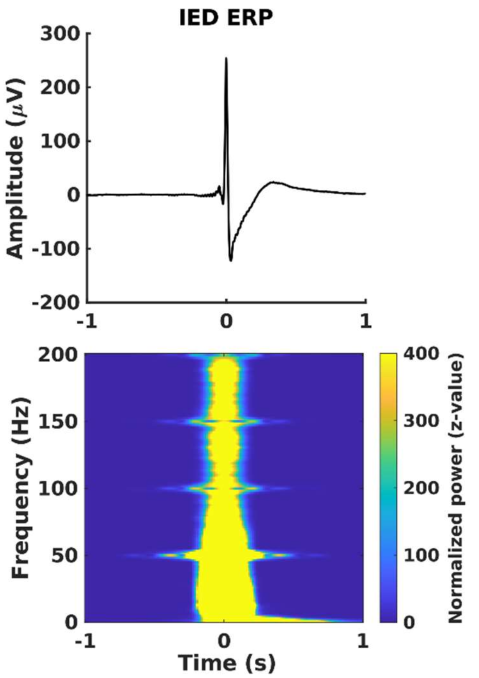


**Figure S2.** *Bottom:* Event related potential (ERP) averaged over all IEDs (1247.19 ± 152.89) detected in the anterior hippocampus shows a phenotypical IED shape with amplitudes ranging from -100 to 300 µV. Filtering sharp transients can yield spurious activity across a broad range of frequencies (see IED TFR in the top panel); therefore, IED periods (1s around the peak) were excluded from event detection and further analysis.

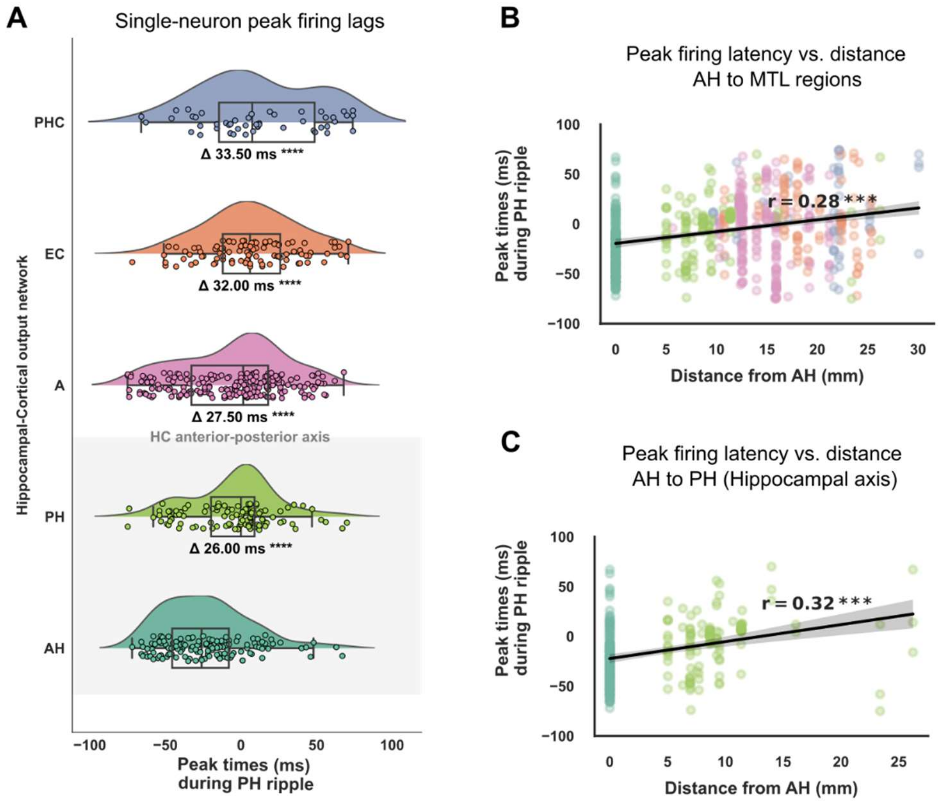


**Figure S3**. (A) Ripple-associated information flow across MTL regions for PH ripple-triggered activity revealed a pattern consistent with AH-triggered dynamics (see Fig. 2). Delta time lags calculated as the difference in median lag between AH and each target region indicate a temporally structured propagation of activity. Along the hippocampal longitudinal axis, PH showed a significant lag of 26 ms relative to AH (Holm-corrected $p = 6.29 \times 10^{-7}$), consistent with posterior progression of activation. Propagation along the hippocampal output axis was also evident: Amygdala (A) exhibited a 27.5 ms lag ($p = 6.29 \times 10^{-7}$), EC exhibited a 32 ms lag ($p = 1.88 \times 10^{-11}$), and PHC a 33.5 ms lag ($p = 1.28 \times 10^{-7}$) all significantly different from AH. All reported values reflect Holm-corrected significance levels. Statistical comparisons were performed using the Wilcoxon rank-sum test on the respective timelag distributions (AH vs. target region). (B) Across medial temporal lobe regions, neurons engaged during PH SWRs demonstrated a significant distance-dependent shift in firing latency, with more distal sites exhibiting progressively later peak activity relative to the PH SWR maximum (Pearson $r = 0.28$, $p = 7.55 \times 10^{-12}$). (C) A similar spatial gradient was evident within the hippocampus itself, where firing latencies increased systematically along the anterior-posterior axis as a function of distance from AH (Pearson $r = 0.32$, $p = 3.19 \times 10^{-7}$). Together, these findings support robust, directional information flow originating in AH and extending to the posterior hippocampus and outward to MTL cortical targets.

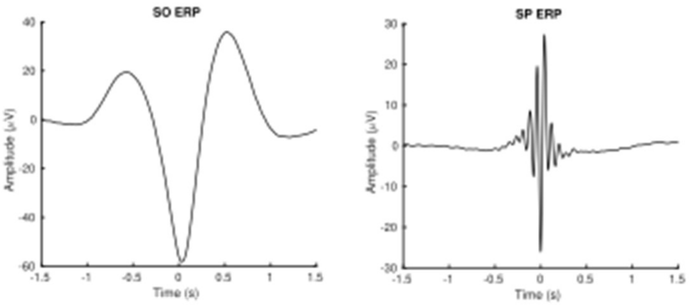


**Figure S4.** ERPs of sEEG-detected (Fz) SOs (1665.43 ± 101.54) and spindles (1937.52 ± 155.06).

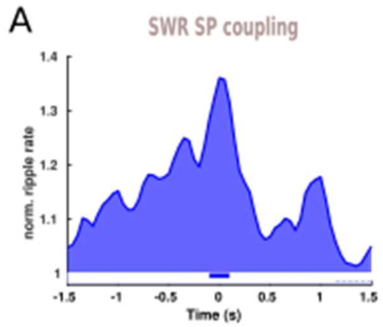


**Figure S5.** SWRs in the anterior hippocampus (AH) show precises temporal coupling to spindles detected at electrode Fz ($p$ = 0.017, –0.1 to 0.15 s; $p$ = 0.002, 1.15 to 1.5 s; two-sided paired-samples t test, corrected across time). PETH shows SWR density centered on the spindle trough.

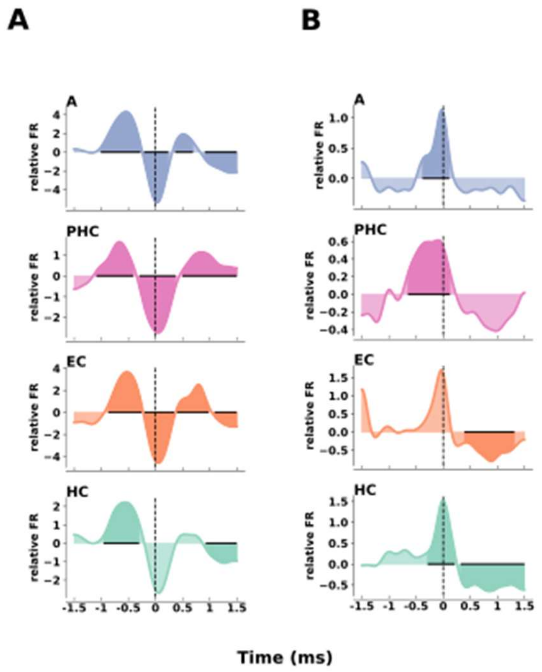


**Figure S6.** Cortical top-down modulation of MUA in t the MTL for all scalp detected SO (**A**) and SP (**B**) events. Shaded areas indicate significant modulations (*p* <0.05, two-sided paired-samples t test, corrected over time, for detailed cluster statistics see Table S3).

**Text S1.** To assess the precision of hippocampal–cortical coordination, we performed time–frequency analysis at frontal EEG Fz during scalp detected SO–SP complexes and SWR-SO–SP triplets (see Fig. 3C–E, top panels). As anticipated, TFR analysis of SO–spindle complex locked to the SO trough revealed robust increases in normalized power within the SO–spindle frequency range (Fig. 3C). This included a significant cluster ($p$ = 0.001, two-sided paired-samples t test, corrected over time and frequency), extending from low frequencies to two spindle-range power increases, precisely timed to the first and second SO up-states. Qualitatively similar enhancements were observed when aligning to spindles nested within SOs, with broadband power elevations extending across the SO–spindle range ($p$ = 0.001; Fig. 3D). Finally, aligning cortical EEG to hippocampal SWRs nested within SO–SP complexes (i.e., SO–SP–SWR triplets) revealed striking hippocampal-cortical synchrony, with significant power increases in the SO–SP range ($p$ = 0.001) precisely time-locked to the SWR peak (t = 0 s), and a secondary cluster in the SO range ($p$ = 0.006; Fig 3E). Notably, this spectral signature closely mirrors the power distribution observed for spindles coupled to SOs. Together, these findings demonstrate temporally precise, spectrally specific coupling between hippocampal SWRs and cortical SO–spindle dynamics, supporting the existence of coordinated mesoscale communication across distant brain regions.
